# Modulation of Filopodial Myosin Function

**DOI:** 10.1101/2025.05.29.656896

**Authors:** Amy Crawford, Jessica K. Schwartz, Emerson Hall, Anne Houdusse, Margaret A. Titus

## Abstract

MyTH-FERM (MF) myosins are essential for filopodia formation in both Metazoa and Amoebozoa (*Dictyostelium* DdMyo7; mammalian Myo10) but their roles in filopodia formation are not fully understood. Taking advantage of a mutation in the highly conserved actin binding interface of another MF myosin, the *jordan* (*jd*) mutation of Myo15A that reduces its function, the impact of altering the myosin-F-actin interaction on filopodia formation was investigated. The *jd* mutation, a D to G substitution (Gong et al, 2022, Sci Adv), was introduced into DdMyo7 or Myo10 and filopodia formation assessed by quantitative analysis of number, length and myosin tip enrichment. The mutation significantly decreased filopodia initiation and tip intensity of both DdMyo7 and Myo10. This is consistent with impairment of myosin-based reorganization of cortical actin at the cortex, with myosins in the initiation complex remaining at the tip as the filopodium extends. The DdMyo7 *jd* mutation did not affect filopodia length. In the case of Myo10, while the *jd* mutation reduced Myo10 velocity of intrafilopodial motility by 40%, resulting in reduced levels of myosin at the filopodia tip, the *jd* mutant did not consistently affect the length of filopodia in different cell lines. This indicates that the major role of Myo10 may not be to promote filopodia elongation by reducing membrane tension, as has been proposed. Rather, the phenotype of the DdMyo7 and Myo10 *jd* mutants indicates that the major role of evolutionarily diverse filopodial myosins is to re-organize actin filaments at the membrane-cortex interface during the critical initiation step.

## Introduction

Cells to move through complex environments powered by continuous extension of membrane protrusions coupled with retraction of the rear of the cell. Protrusions such as lamellipodia and filopodia are driven by ongoing actin polymerization against the plasma membrane and they not only extend the cell front, but they also make strong adhesions with the extracellular environment. Filopodia are slender membrane protrusions filled with parallel bundles of actin that cells use to both explore and interact with their environments. Filopodia tips harbor components of the cellular adhesion machinery, such as integrins and talin, enabling them to make first contact with the extracellular environment. Following that initial interaction, filopodia adhesions can mature into strong adhesion sites, or generate pulling forces that enable the cell to organize the extracellular matrix to facilitate migration or bring captured bacterial prey into the cell. Filopodia are made by a diverse range of cells throughout phylogeny, indicating that they are ancient cytoskeletal structures (1).

The mechanism of filopodia formation is not fully understood. Initiation of a filopodium is promoted by local activation of a small GTPase at the plasma membrane that recruits actin regulators, notably VASP or a formin, and a myosin to assemble an initiation site and propel elongation of the filopodium. Filopodial myosins are members of the MyTH-FERM (MF; **my**osin **t**ail **h**omology 4 - band **4**.1, **e**zrin, **r**adixin, **m**oesin) family characterized by the presence of their namesake domain at the end of the C-terminal tail. The MF domain is a protein interaction module and is known to interact with the C-terminal tail of receptors, such as β-integrin, and microtubules, for example. MF myosins are localized to the tips of membrane extensions supported by parallel bundles of actin, namely filopodia (amoeboid DdMyo7, Myo10), microvilli (Myo7B) and stereocilia (SC) (Myo7A, Myo15A), and are associated with their growth, organization and maintenance (2, 3).

Metazoan Myo10 and amoeboid DdMyo7 are evolutionarily distant MF myosins essential for filopodia formation in mammalian cells and amoebae, respectively (4, 5). They are recruited to the plasma membrane where they concentrate at initiation sites and then remain at the tip as a filopodium elongates (6, 7). DdMyo7 and Myo10 are functional homologues, but while they share some features in their motor properties, they are recruited to the cortex or membrane differently. Both motors exist in an autoinhibited state in the cytosol and upon recruitment to the cortex they dimerize (8, 9). Forced dimers of the motor domain of DdMyo7 and Myo10 can promote filopodia formation, but not as efficiently as the full length myosin, indicating that the tail domain is important for filopodial myosin function (8, 10, 11). However, Myo10 is an anti-parallel dimer and a fast, processive motor that moves optimally along bundled actin filaments (660 nm/sec) (12). In contrast, DdMyo7 is a much slower processive motor that does not move significantly faster on bundled actin - 40 nm/sec for a single filament vs 56 nm/second on fascin-bundled actin (13). It is not yet known if DdMyo7 and Myo10 play similar or different roles during filopodia initiation and extension.

Several roles have been proposed for filopodial myosins in initiation and extension. One model proposes that the dimeric myosin organizes the ends of growing actin filaments into parallel bundles at the membrane-cortex interface, likely in collaboration with the actin polymerase/bundle VASP, and then transports Ena/VASP along the filopodial actin bundle to promote ongoing growth (14). Support comes from the finding that a forced dimer of a filopodial myosin, either Myo10 or DdMyo7, can promote filopodia formation (8, 11) and VASP is seen to co-transport with Myo10 along the length of a filopodium (15, 16). However, recent work has shown that tethering of a monomeric Myo10 myosin motor to the membrane results in tip accumulation and can promotes filopodia extension (17). Similarly, chimeric myosins generated by exchanging different myosin motors into Myo3A that binds both the actin core of SC as well as the membrane, can also drive filopodia elongation (18). The results from these studies suggest that the tip-localized motors could generate a downward force on actin filaments that relieves membrane tension, allowing for the ongoing addition of actin monomers. Interestingly, this activity is only seen for myosin motors with roles in filopodia or stereocilia extension, Myo3A (SC), Myo10 (filopodia) and Myo15A (SC) (17–19), indicating that there could be a highly specific mode of interaction between actin and the motor at the tips of these specialized protrusions that promotes actin polymerization against the membrane.

Myo15A, is an MF myosin essential for full elongation of SC that has been proposed to promote actin polymerization at the SC tip. Mutation of a key residue in the Myo15A motor, D1647G (*jordan* mutation, *jd*) causes progressive hearing loss due to impaired growth of stereocilia (20, 21). This mutation is of particular interest as it modulates the functionality of the Myo15A motor and does not abolish motor function altogether. In contrast to the canonical *sh^2^* mutation where mutant Myo15A and its elongation complex (EC) cargo (WRN, EPS8) are not localized to the SC tip, the *jd* mutant and the EC remain at SC tips, although their overall levels are moderately decreased (22, 23). In spite of the presence of the WRN and EPS8 at the tip, the SC do not elongate, suggesting dysregulation in actin polymerization.

The *jd* mutation is in a highly conserved region of the motor that makes initial contact with actin, the helix turn helix (HTH) (Fig 1A,B), a region of the actomyosin interface where the motor bridges two actin monomers (20). Surprisingly, the Myo15A motor can promote nucleation of actin in vitro and this activity is disrupted by the *jd* mutation (20, 21). The effect of the *jd* mutation on SC elongation and actin nucleation in vitro led to the proposal that the role of Myo15A is SC elongation is to directly promote actin polymerization at the growing tips of SC in collaboration with the EC. The proposed role for Myo15A in promoting elongation of parallel bundles of actin by actin polymerization activity in SC raised the question of whether MF myosins might have similar roles in filopodia. The HTH is a conserved region of the motor in the lower 50K region involved in motor binding to F-actin (24, 25) and the *jd* residue is a highly conserved amino acid in that region (Fig 1A,B). The majority of myosins have an E in that position, including both the DdMyo7 and Myo10 motors, while Myo15A has a conservative D substitution. Taking advantage of the high sequence similarity in the HTH region of myosins, the impact of a ‘*jd*’ mutation in the HTH region on filopodia formation was tested. Making a single missense mutation that modulates myosin motor function in a critical region of the actomyosin interface provides a way to precisely probe the role of myosins in their cellular context, a first step towards understanding whether the myosins involved in elongation of filopodia and stereocilia have shared mechanisms.

**Figure 1.**
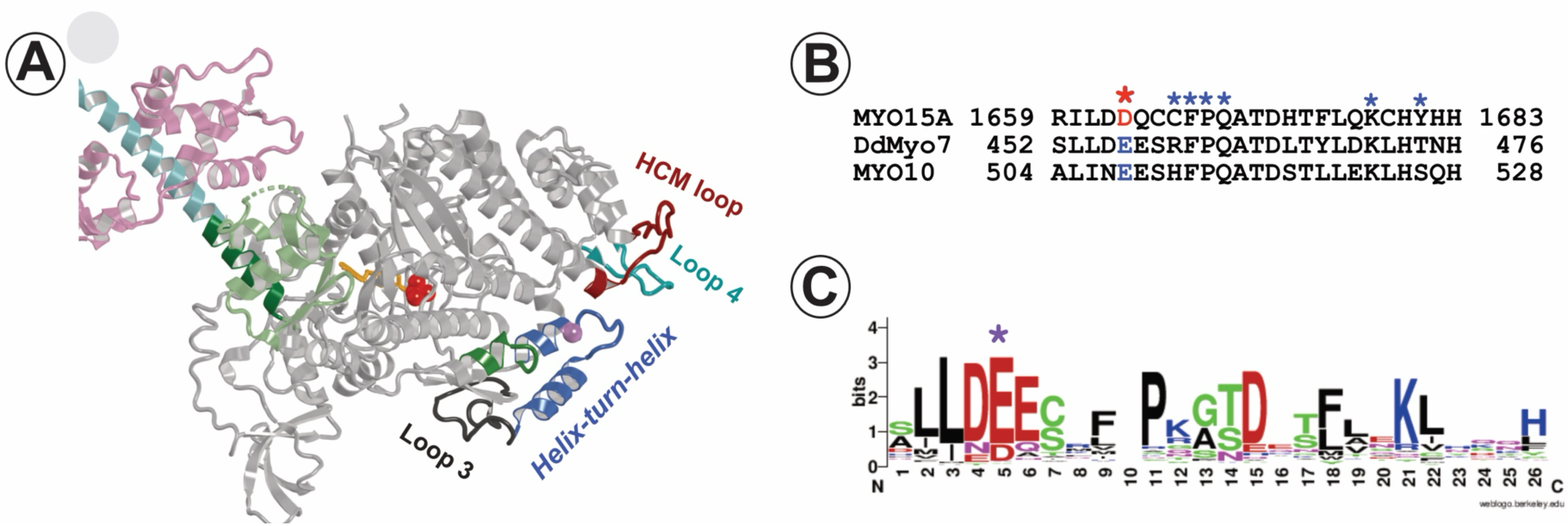
The actin-binding HTH region of the myosin motor domain. **A.** Homology model of the DdMyo7 motor domain based on the Myo10 motor in the pre-power stroke state (ADP-Pi; PDB 510H) showing the helix-turn-helix (HTH) on the lower 50K region of the motor (blue); along with the HCM loop (maroon), Loop 4 (aqua) and Loop 3 (black) regions important for actin binding. The red balls and orange stick represent bound nucleotide, the green helix is the activation loop and the purple ball on the HTH is the ‘*jd*’ residue at the end of the first helix. **B.** Alignment of the HTH sequences of Myo15A, DdMyo7 and Myo10. red * denotes the ‘*jd*’ residue. * actin interacting residues, **C.** LOGO representation of alignment of the HTH region from 136 different myosin motor domains from 53 different classes plus two divergent orphan myosins across phylogeny. Note that the gap at position 10 is due to the presence of a highly variable or no amino acid at that position. The red * indicates the ‘*jd*’ residue and the blue * indicate residues on the actin interface with actin.

## Results

Alignment of the HTH sequence from representative myosin family members from 53 different classes across a wide number of species shows strong conservation (FIG 1B,C). The Myo15A *jd* residue, D1647 of the motor domain, is in the position of one of the most highly conserved residues. However, the majority of myosins, including the two evolutionarily distant filopodial myosins, DdMyo7 and mammalian Myo10 have an E instead of a D (79% E, 15% D) (Fig 1C). Both of these residues have the same negative charged with the side chain differing by one carbon, a relatively modest difference. The impact of mutating the ‘*jd*’ residue on the function of these two filopodial myosins was assay using live cell imaging and quantitative image analysis of filopodia formation (see Methods; (26)). Several parameters were measured to pinpoint which aspect of myosin function in vivo is impacted by the mutation - 1) The fraction of cells making filopodia indicates overall myosin functionality, 2) the cortex to cell body ratio is a measure of cortical recruitment of DdMyo7 that requires relief of autoinhibition and actin dynamics or membrane recruitment of Myo10 by PI(3,4,5)P3 generation, 3) filopodia count provides a measure of the myosin’s ability to generate initiation foci through reorganization of the actin network adjacent to the membrane that result in filopodia formation, 4) filopodia length assesses the elongation machinery and 5) the tip to cell body ratio is an indirect measure of size of the initiation site, as it has been shown that once an initiation foci forms at the cortex, it becomes the filopodia tip and the intensity remains constant throughout elongation (7, 8, 27). For Myo10, tip intensity also increases as the filopodium elongates, due to intrafilopodial motility of the motor from the filopodia to the tip that may contribute to sustained growth (17, 28). Together, determining how these different parameters of filopodia formation are altered provides a comprehensive understanding of filopodial myosin function is perturbed by a motor domain mutation.

### Impact of the jd mutation on DdMyo7-based filopodia formation

Two distinct mutations were introduced into the DdMyo7 motor of full-length mNeon-DdMyo7 (referred to hereafter as DdMyo7), one is the equivalent *jd* mutation (E456G) and the other a conservative mutation to D, the WT residue in the Myo15A motor (E456D). The DdMyo7-E456G mutant exhibited significantly reduced efficiency of filopodia formation, with only 28% of E456G cells making filopodia as compared to 77% for the control wild type (WT) DdMyo7 (Fig 2A, Table 1). The number of filopodia/cell was also substantially reduced for both mutants (WT: 4.38 ± 0.169, E456G 1.60 ± 0.057 p < 0.0001) (Fig 2B, Table 1). The decrease in filopodia formation is correlated with a loss of cortical enrichment (WT: 1.13 ± 0.008, E456G 1.02 ± 0.003 p < 0.0001) (Fig 2C, Table 1). A lower number of filopodia in the DdMyo7-E456G cells compared to WT indicates that there must be fewer initiation events that lead the formation of a filopodium (Fig 2, Table 1). Interestingly, the conservative E to D substitution also results in reduced filopodia formation. Only 41% of the cells expressing DdMyo7-E456D make filopodia and these have 2.13 ± 0.088 filopodia/cell, a significant drop in filopodia number compared to WT (p< 0.0001) but notably more than the E456G mutant (Fig 2A,B, Table 1). Cortical enrichment is also reduced (1.07 ± 0.005) but not as much as seen for the E456G mutation (Fig 2C, Table 1). Thus, either a significant (E to G) or minor (E to D) change in the highly conserved HTH amino acid impacts the ability of DdMyo7 to promote filopodia formation. DdMyo7 is recruited to a dynamic actin-rich cortex that depends on VASP-mediated actin polymerization via its motor domain (27). The predicted weakening of the actomyosin interaction due mutation of E456 in the DdMyo7 HTH region is consistent with the observed reduction in cortical targeting and decreased filopodia formation.

**Figure 2.**
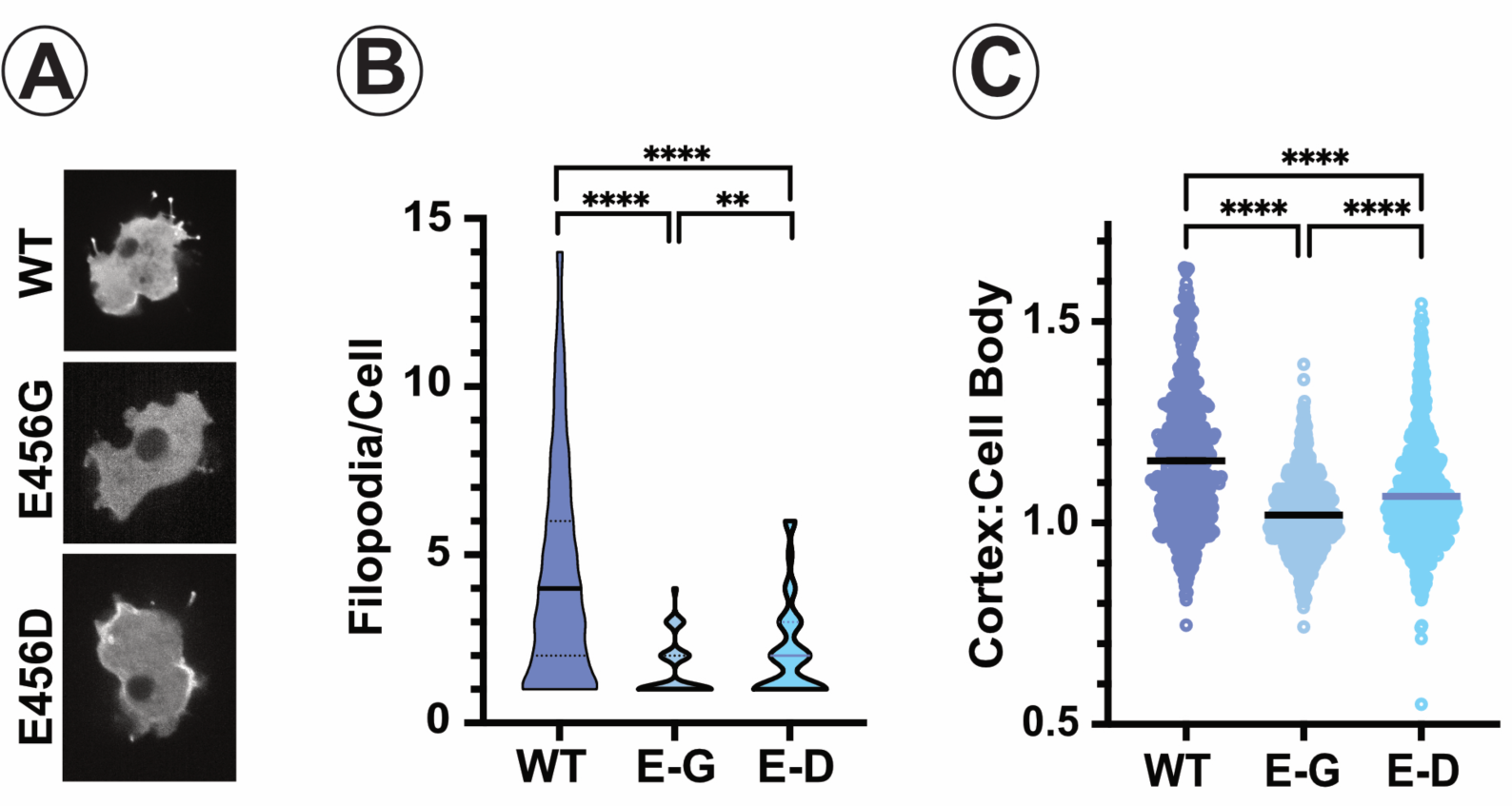
Mutation of the DdMyo7 ‘*jd*’ residue significantly inhibits filopodia formation. Quantitative analysis of filopodia formation rescue by mNeon-full-length wild type (WT), E456G (E-G) or E456D (E-D) DdMyo7-expressing *myo7* null cells using filoTips. **A.** Representative confocal micrographs of each cell line that show differences in the number of bright filopodia tips and cortical enrichment in *myo7* null cells expressing the HTH mutant myosins. **B.** Filopodia per cell. The mean is indicated with a thick bar on the plot for each line and dashed lines indicate the quartiles. **C.** Cell body to cortex ratio. The mean is indicated with a thick bar on the plot. Shown are the results from the analysis of two independent cell lines for each DdMyo7. Two independent cell lines were analyzed for each mutant and a minimum total of four independent assays were performed for each mutant. ** p < 0.03 **** p < 0.0001

**Table 1.** Impact of HTH mutants on DdMyo7 filopodial function.

|  | **WT** | **E546G** | **E546D** | **I426A** |
| --- | --- | --- | --- | --- |
| cells with filopodia | 77 %<br>(379/495) | 28%<br>(217/770) | 41%<br>(293/723) | 54%<br>(448/831) |
| cortex:cytoplasm ratio ± SEM | 1.13 ± 0.008 | 1.02 ± 0.003**** | 1.07 ± 0.005 **** | 1.07 ± 0.004 **** |
| n cells | 494 | 766 | 716 | 831 |
| filopodia number ± SEM | 4.38 ± 0.169 | 1.60 ± 0.057**** | 2.13 ± 0.088**** (°) | 2.89 ± 0.105**** |
| n cells | 367 | 207 | 266 | 426 |
| filopodia length ± SEM | 1.57 ± 0.025 | 1.30 ± 0.041 | 1.41 ± 0.033**** (^) | 1.27 ± 0.022**** |
| n filopodia | 650 | 163 | 217 | 471 |
| tip intensity ± SEM | 1.71 ± 0.024 | 0.73 ± 0.016**** | 1.07 ± 0.022**** | 1.46 ± 0.022**** |
| n measured tip intensity | 652 | 168 | 220 | 471 |
| #, N | 5, 495 | 4, 770 | 4, 723 | 4,831 |

Summary of filopodial formation parameters for wild type and mutant DdMyo7 expressed in the *myo7* null line. The value for filopodia/cell does not include cells that did not make any filopodia.

Models for myosin-based filopodia formation propose that motor activity at the filopodia tip is critical either for promoting actin polymerization or reducing membrane tension at the tips which would facilitate addition of G-actin monomers (17, 19, 21). In the case of Myo15A, the tallest row of stereocilia in the cochlear inner and outer hair cells of *Myo15a^jd/jd^* mice are about 50% shorter than those of *Myo15a^+/+^* mice (21). If DdMyo7 plays a critical role in promoting actin polymerization at the filopodia tip, then mutation of the *jd* residue would be expected to have a significant effect on filopodia length. Filopodia length is decreased in both the E456G and E456D mutants, but only to a modest extent (WT: 1.57 ± 0.025 µm, E456G: 1.30 ± 0.041 µm, 20% shorter; E456D: 1.41 ± 0.033 µm, 10% shorter; with p> 0.0001 for each mutant compared to WT) (Fig 3A, Table 1). These results show that DdMyo7 does not play a significant role in filopodia elongation.

**Figure 3.**
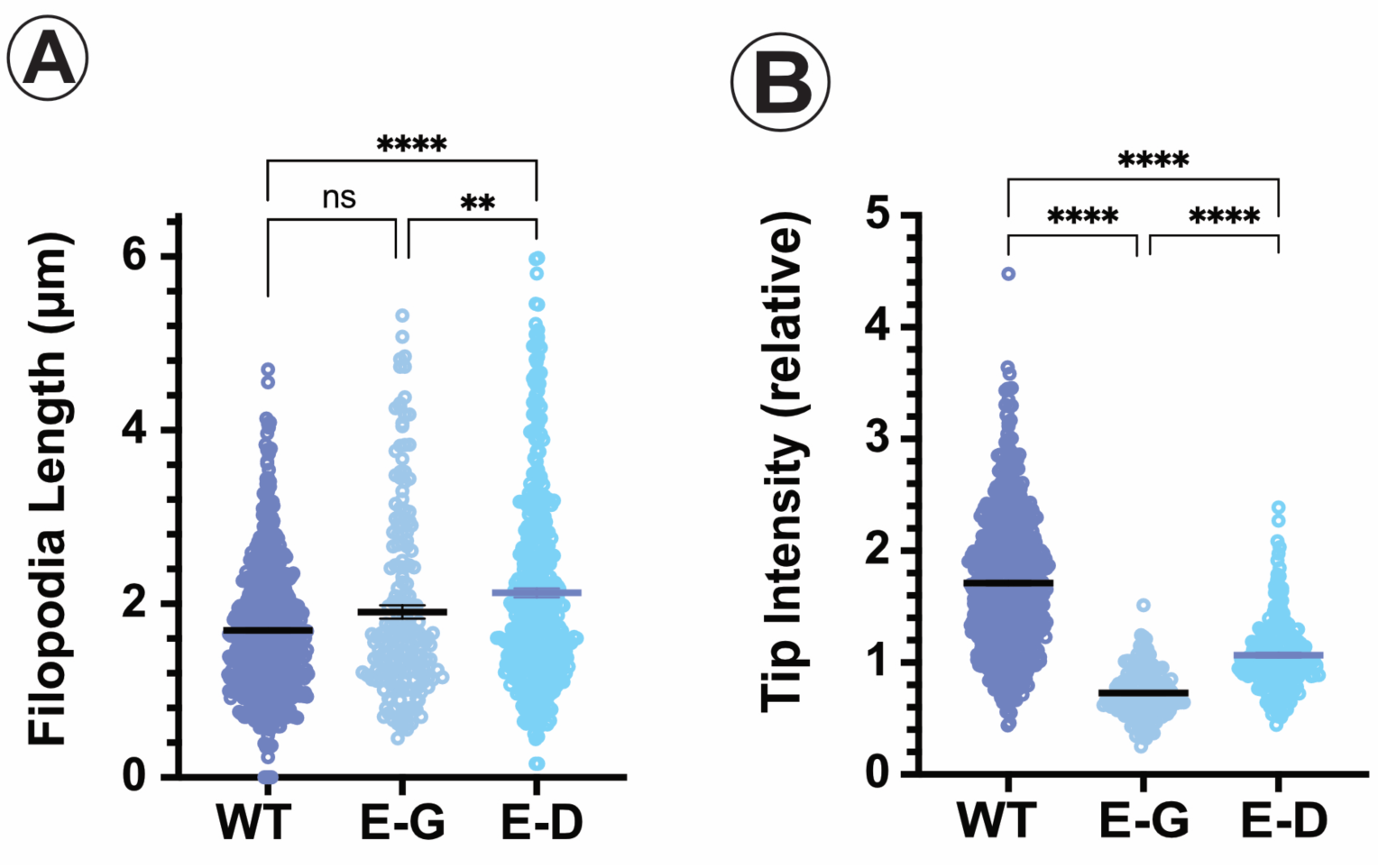
The DdMyo7 ‘*jd*’ mutation does not impact filopodia length. Quantitative analysis of filopodia formation in wild type cells expressing either the wild type (WT), E456G (E-G) or E456D (E-D) mNeon-DdMyo7 using filoTips. **A.** Filopodia length. The mean is indicated with a thick bar on the plot for each line and thinner lines indicate the SEM. **B.** Filopodia tip intensity. The mean is indicated with a thick bar on the plot for each line. Shown are the results from the analysis of two independent cell lines for each DdMyo7, Two independent cell lines were analyzed and a minimum of four independent assays were performed for each mutant. ** p < 0.002 **** p < 0.0001, ns = not significant.

The slightly shorter filopodia observed in the the DdMyo7-E456G expressing cells have significantly dimmer filopodia tips, ∼ 45% lower tip intensity compared to WT (WT: 1.84 ± 0.04, E456G: 0.86 ± 0.02 tip/cell body intensity ratio, p < 0.0001) and DdMyo7-E456D tips are ∼ 40% dimmer (1.15 ± 0.02 tip/cell body intensity ratio, p < 0.0001) (Fig 3B, Table 1). The reduced intensity indicates that fewer motors are present at the tip and this could reflect less efficient formation of initiation foci, as fewer motors are localized at the cortex and less are available for recruitment into an initiation site. It cannot be attributed to a decrease in DdMyo7 intrafilopodial motility as this has not been observed and the slow speed of the motor (∼ 40 nm/sec) is likely lower than retrograde flow (7, 13).

Taken together, characterization of the DdMyo7-E456G and -E456D phenotypes indicates that DdMyo7 motor activity and force generation is critical for the initiation step of filopodia formation and not for filopodia extension.

### Impact of the jd mutation on Myo10-based filopodia formation

WT or mutant Myo10 was expressed in COS-7, U2-OS or HeLa cells to determine how a change in the HTH actin-binding interface might affect its role in filopodia formation. It should be noted that these three cell lines natively make varying numbers of filopodia. COS-7 cells make few, if any, filopodia (4), U2-OS cells make a modest number (28) and HeLa cells make significant numbers of filopodia.

Expression of WT Myo10 in COS-7 cells results in robust production of filopodia but the Myo10-E508G mutant generated only 20% the number of filopodia as seen for WT Myo10 (68.21± 5.26 vs 13.79 ± 1.73, p < 0.0001) (Fig 4A,B, Table 2). Unlike the DdMyo7-E456G mutant, the cortex/cell body ratio of the Myo10-E508G mutant is similar to that of WT Myo10. This is expected as the mode of recruitment for the two myosins differs (9, 27). Myo10 is recruited to the cortex occurs via PI(3,4,5)P3 production (29), while DdMyo7 requires actin binding for its targeting and activation (27). The overall number of filopodia is reduced to 40.15 ± 4.16 filopodia/cell in the Myo10-E508D mutant, significantly lower than WT but not as much as seen for Myo10-E508G, and no change is observed in the cortex/cell body ratio (Fig 4, Table 2). Similar reductions in filopodia numbers are seen when the Myo10-E508G and -E508D mutants are expressed in U2-OS and HeLa cells, demonstrating that the effect on filopodia formation is not cell-type specific (Suppl Table 1). These results show that, as seen for DdMyo7, mutation of the *jd* residue significantly impacts Myo10-mediated production of filopodia. This is likely due to a weakened actomyosin interaction that impairs the ability of the motor to contribute to the re-organization of cortical actin filaments needed for initiation. Furthermore, it also reveals that two different filopodial myosins show the same extent of reduced activity when the *jd* mutation or a conservative substitution is introduced into the HTH.

**Figure 4.**
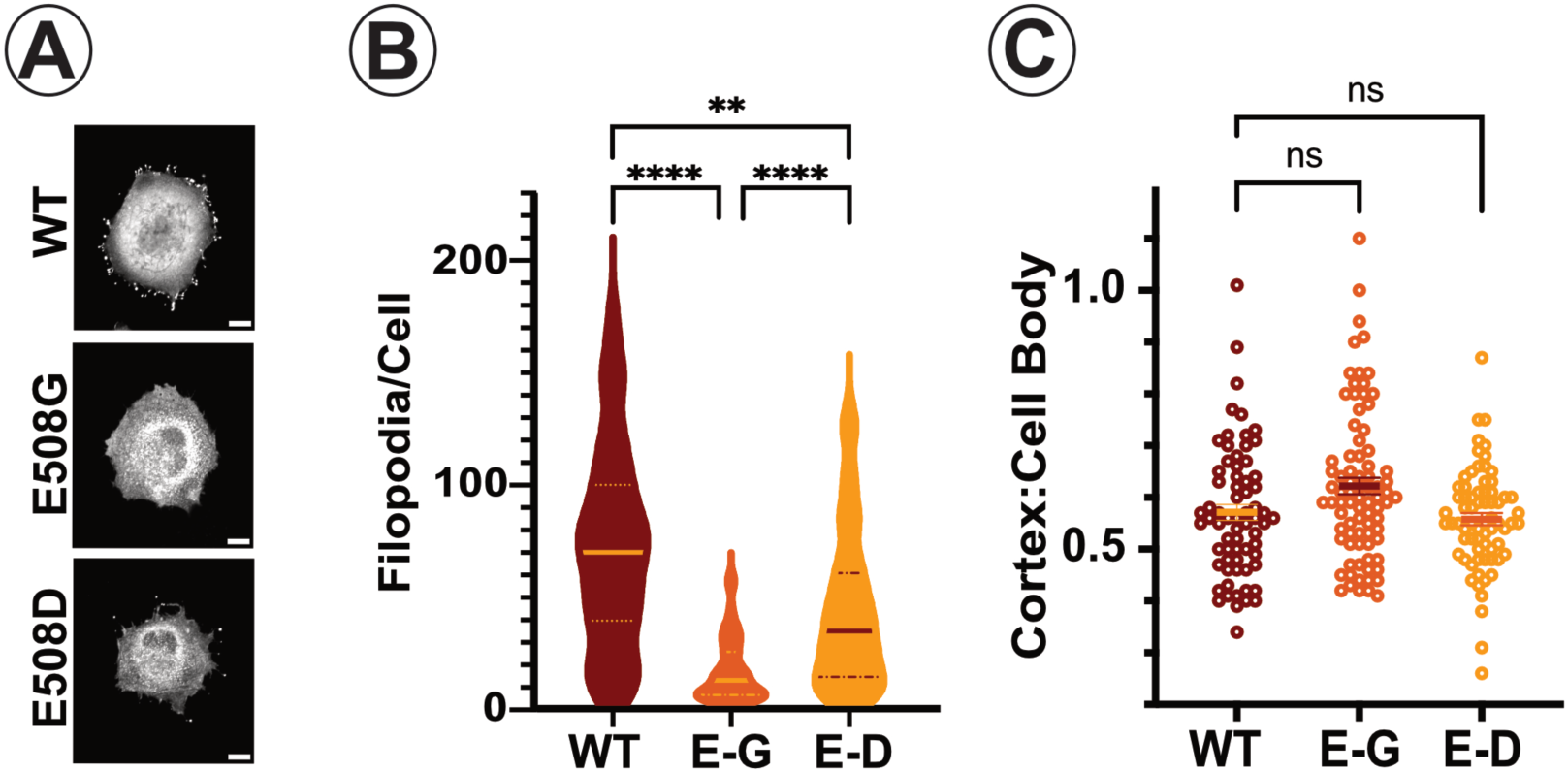
Mutation of the Myo10 ‘*jd*’ residue impairs filopodia formation. Quantitative analysis of filopodia formation by full-length eGFP-wild type (WT), E508G (E-G) or E508D (E-D) Myo10-expressing COS-7 cells using filoTips. **A.** Representative images of COS-7 cells expressing wild type and mutant Myo10. Scale bar = 10 µm. **B.** Filopodia per cell. The mean is indicated with a thick bar on the plot, smaller dashed lines indicate quartiles. **C.** Cortex:cell body ratio of Myo10 localization. The mean is indicated with a thick bar on the plot. Shown are the results from the analysis of two independent cell lines for each transfection. Two independent cell lines were analyzed for each mutant and a minimum total of four independent assays were performed for each mutant. * p < 0.004 ** p < 0.03 **** p < 0.0001

**Table 2.**
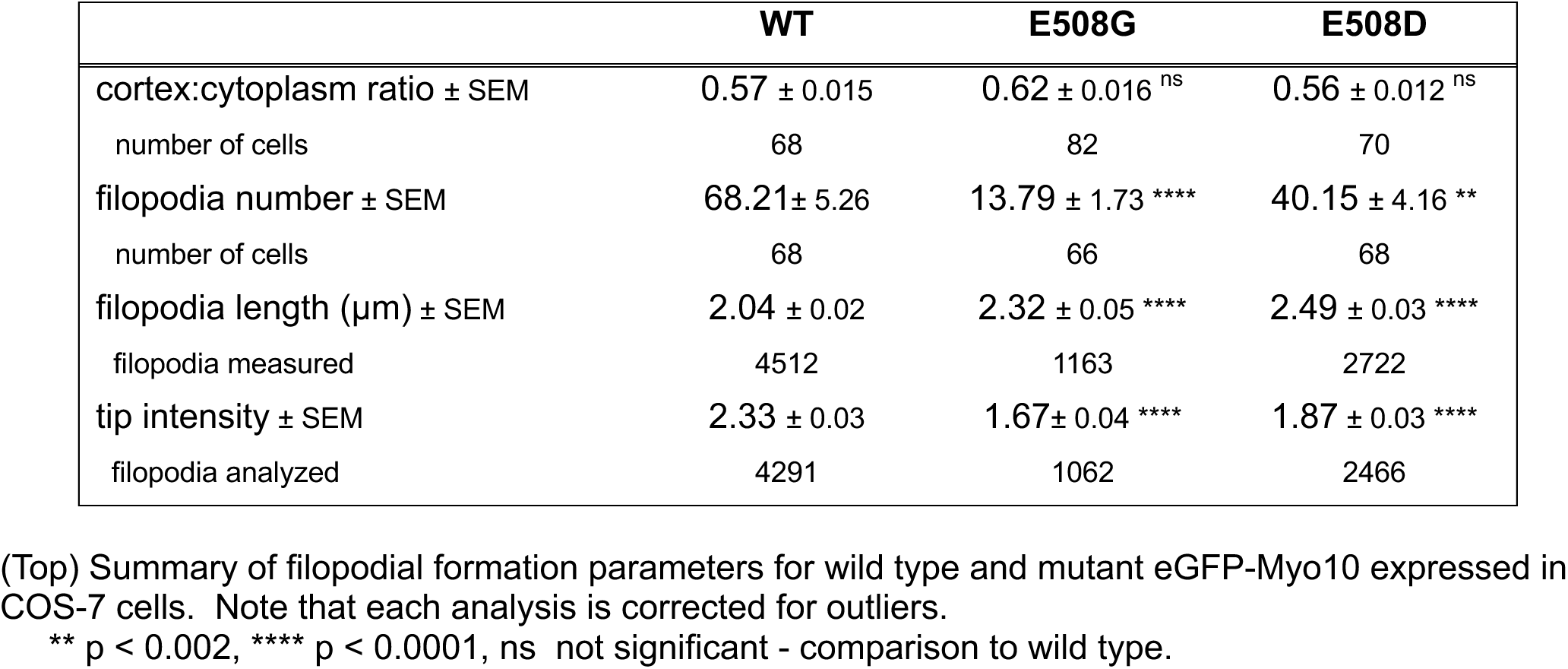
Impact of HTH mutants on Myo10 filopodial function in COS-7 cells.

Intrafilopodial movement of Myo10 from the filopodia tip to the base results in accumulation of Myo10 to filopodia tips (30) and has been suggested to play a role in elongation (17, 28). TIRF imaging was used to measure the intrafilopodial motility of the full-length WT and mutant Myo10s in HeLa cells (30) to determine the effect of the *jd* mutations on in vivo motor activity. HeLa cells were imaged within 6 - 8 hr of transfection to ensure a modest level of expression that allowed for visualization of eGFP-Myo10. Motile green punctae were clearly visible within the filopodia shaft, with most of them moving towards the tipin all three cell lines (Fig 5). The mean velocity of eGFP-Myo10 punctae was 601 ± 25 nm/sec, similar to previously reported for this myosin (30). The Myo10-E508G and Myo10-E508G moved significantly slower (p < 0.0001), 411± 11 nm/sec and 498 ± 14 nm/sec (Fig 5, Table 3), respectively, as would be expected based on the observed reduction in the gliding motility seen for the Myo15A *jd* myosin motor (21). While the 40% slower velocity observed for Myo10-E508G is not as substantially reduced as seen for the Myo15A *jd* motor domain in gliding motility assays (55% reduction) (21), but overall the impact of this mutation is quite significant for both myosins. These results show that mutation of the *jd* residue decreases Myo10 motor transport function within filopodia. The length of filopodia in COS-7 cells expressing WT or mutant Myo10s was measured to determine the impact of impairing Myo10 motor function on filopodia elongation. Unexpectedly, the filopodia in Myo10-E508G and -E508D expressing cells were longer than the WT control (WT: 2.04 ±0.02 µm, E508G: 2.32 ± 0.05 µm, E508D: 2.49 ± 0.05 µm, p < 0.0001 comparison between WT and mutants) (Fig 6A, Table 2). The tip intensity was, however, reduced for both mutants (WT: 2.33 ± 0.03, E508G: 1.67 ± 0.04, E508D: 1.87 ± 0.03, p < 0.0001 comparison between WT and mutants) (Fig 6B, Table 2). This contrasts with both U2-OS and HeLa cells where a decrease in both filopodia length and tip intensity is seen (Supp Fig 1, Supp Table 1). The filopodia of Myo10-E508G expressing HeLa cells are almost 50% shorter than those expressing Myo10-WT (WT: 4.71 ± 0.04 µm, E508G: 2.52 ± 0.03 µm, p < 0.0001) while the Myo10-E508D filopodia length is unchanged (E508D: 4.84 ± 0.05 µm). For both mutants, there is a significant reduction in tip intensity, especially in the Myo10-E508G cells (WT: 2.39 ± 0.02, E508G: 1.24± 0.02, E508D: 1.75 ± 0.02, p < 0.0001 comparison to WT) (Suppl Fig 2, Supp Table 1). In the case of U2-OS cells, a modest decrease in length is observed for Myo10-E508G expressing cells, they are ∼20% shorter (WT: 2.79 ± 0.03, E508G: 2.33 ± 0.03, p <0.0001), while the Myo10-E508D filopodia are slightly longer than the WT filopodia (2.91 ± 0.03, p < 0.0001) (Supp Fig 1, Supp Table 1). Similar to what is observed in COS-7 and HeLa cells, the tip intensity of mutant expressing cells is reduced compared to the WT control (WT: 2.61 ± 0.03, E508G: 1.82 ± 0.03, E508D: 2.22 ± 0.03, p < 0.0001 comparison to WT) (Supp Fig 2, Supp Table 1). Together, these results show that introduction of the *jd* mutation (E508G) into Myo10 can decrease filopodia length, depending on the cell line. It appears that the greater the average length of WT filopodia extended by a given cell, the greater the impact. HeLa cells extend the longest filopodia of the three cell lines, ∼ 4.7 µm vs ∼ 2.8 µm and ∼ 2 µm for U2-OS and COS-7, respectively. The Myo10-E508G mutant has the most significant effect on the filopodial length in these cells - reducing the length by 50% while the Myo10-E508D mutation does not affect filopodial length in any cell line. However, both the E508G and E508D mutations do decrease tip intensity, with the E508G mutation resulting in a greater decrease than the E508D. The varied impact of the *jd* mutations on filopodia length and the finding that mutations such as E508D mutation does not reduce filopodia length but does diminish the amount of Myo10 at tip, the results indicate that Myo10 is not fundamentally important for filopodia elongation. While it may contribute to reduction of membrane tension at the tip that would facilitate extension, this potential function does not appear to be essential. Instead, the overall results are consistent with a major role for Myo10 in initiation, similar to what is found for DdMyo7.

**Figure 5.**
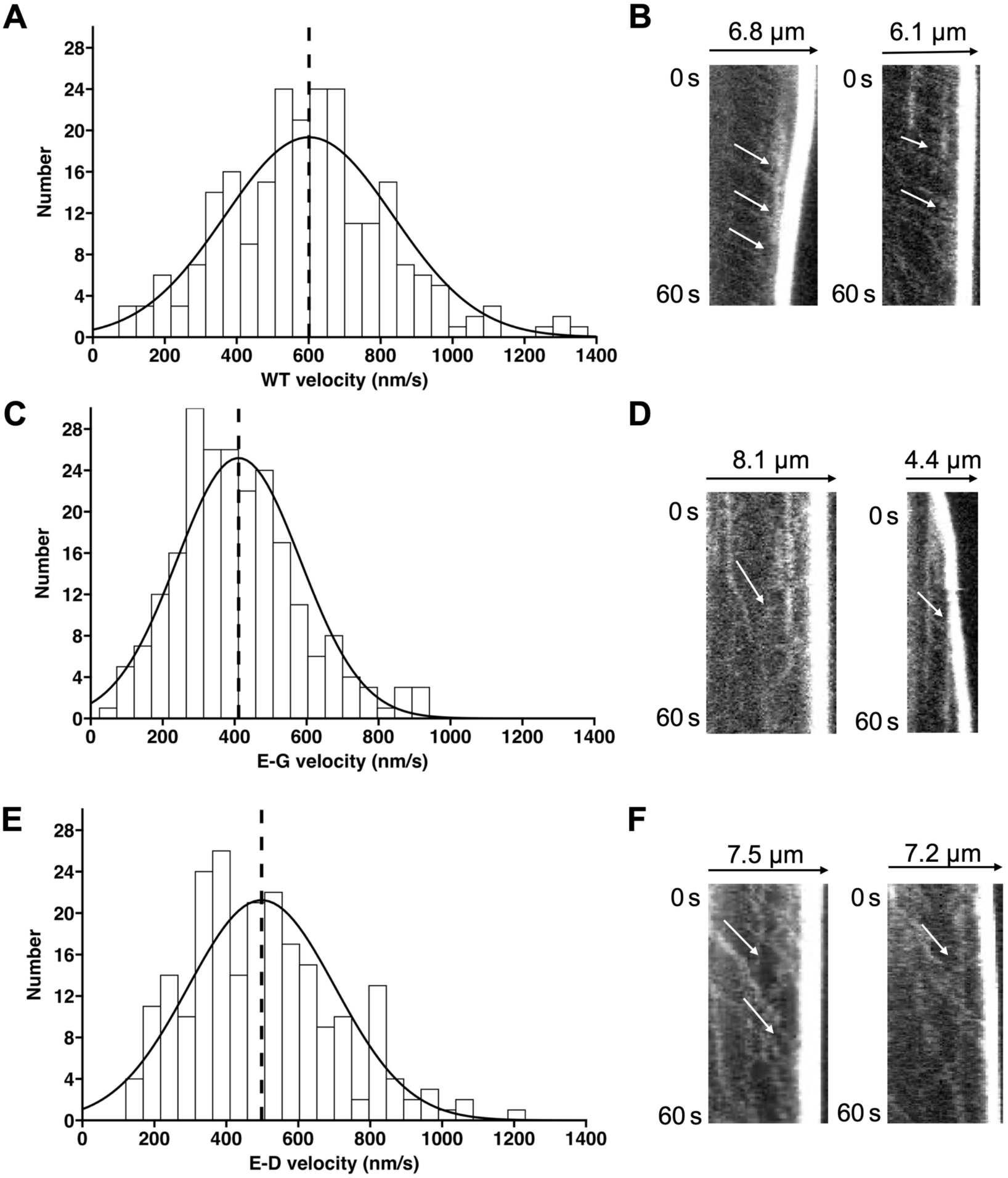
Mutation of the Myo10 ‘*jd*’ residue reduces intrafilopodial motility. Intrafilopodial motility of eGFP-Myo10 punctae of full-length wild type (WT), E508G (E-G) or E508D (E-D), in Hela cell filopodia measured using TIRF microscopy **A, C, E.** Histogram showing the measured velocities for WT, E508G and E508D, respectively. Data fit to a Gaussian curve, and dashed line indicates the mean velocity, **B**,**D,F.** Representative kymographs of movement of wild type (WT), E508G (E-G) or E508D (E-D) punctae moving from filopodia base (left side) to tip (right side).

**Figure 6.**
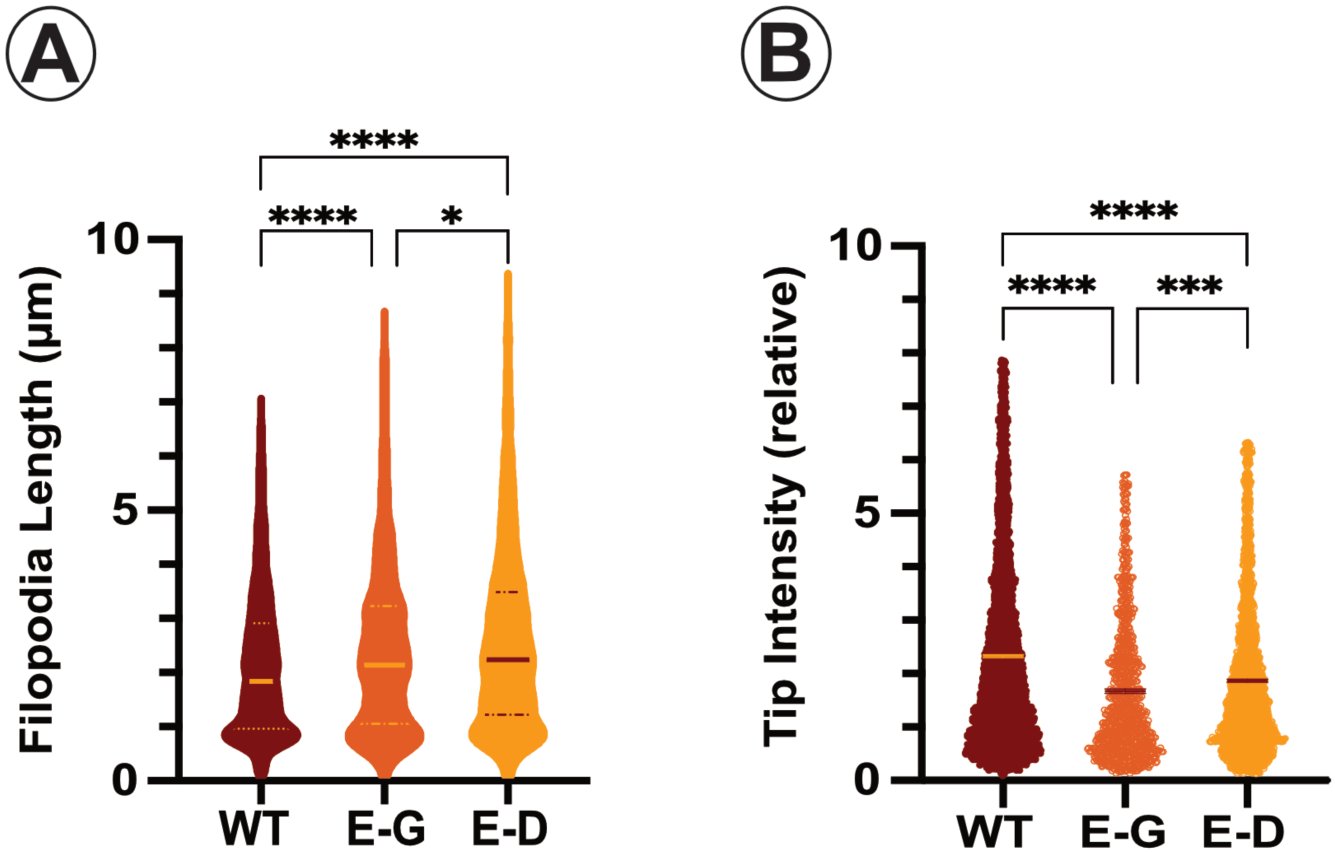
The Myo10 ‘*jd*’ mutation does not impact filopodia length in COS-7 cells. Quantitative analysis of filopodia formation in wild type cells expressing either the wild type eGFP-Myo10 (WT), E508G (E-G) or E508D (E-D) using filoTips. **A.** Filopodia length. The mean is indicated with a thick bar on the plot for each line and thinner lines indicate the quartiles **B.** Filopodia tip intensity. The mean is indicated with a thick bar on the plot for each line. Shown are the results from the analysis of two independent cell lines for each myosin. Two independent cell lines were analyzed and a minimum of four independent assays were performed for each mutant. * p < 0.02, *** p < 0.005, **** p < 0.0001

**Table 3.** HTH mutations reduce Myo10 intrafilopodial motility.

|  | **WT** | **E508G** | **E508D** |
| --- | --- | --- | --- |
| mean velocity ± SEM, nm/sec | 601 ± 15 | 411 ± 11**** | 498 ± 14**** |
| # of experiments | 2 | 2 | 2 |
| no. cells | 25 | 49 | 38 |
| filopodia analyzed | 74 | 94 | 83 |
| measurements | 235 | 225 | 225 |

Summary of intrafilopodial motility from filopodial base to tip for wild type and mutant eGFP-Myo10 expressed in HeLa cells.
\*\*\*\* $p < 0.0001$ , comparison to wild type velocity.

## Discussion

The initial interaction of the myosin motor with F-actin is mediated by the conserved HTH region (Fig 1) (24, 25). Myosin-mediated filopodia formation requires force generation during the earliest stages of initiation, aligning and bundling cortical actin filaments at the interface of the membrane and actin cortex. Filopodial myosins are also proposed to be important for tip extension, where membrane-associated motors may generate a downward force to relieve membrane tension, permitting addition of G-actin monomers (17, 18). In the case of Myo10, it motors up the filopodia shaft, either supplying actin polymerization proteins or contributing to the pool of tip-associated myosins that could be important for reducing membrane tension (17, 30). More recent work has suggested that a myosin motor domain may directly play a role in actin polymerization at the tip of membrane protrusions supported by parallel bundles of actin, based on the in vivo and in vitro phenotypes of a mouse Myo15A mutant, *jd*, that impairs SC elongation (20, 21). Disruption of the HTH-F-actin interaction is predicted to affect any one or all of these aspects of myosin-driven filopodia formation. The *jd* mutation lies in the HTH and it modulates but does not eliminate myosin motor activity. Taking advantage of this finding, the mutation was introduced into the equivalent HTH ‘*jd*’ amino acid in two evolutionarily distant filopodial myosins to gain insight into how myosin-based force generation contributes to filopodia formation.

The E to G mutation in the HTH of both DdMyo7 and Myo10 decreases filopodia production (Figs 2B, 4B, Tables 1, 2, Supp Fig 1, Supp Table 1). DdMyo7-E456G expressing cells generate 65% fewer filopodia and Myo10-508G cells also make fewer filopodia, from 60% (HeLa cells) to 50% (COS-7, U2-OS) less. In addition to lower numbers of filopodia, filopodia tip intensity is also reduced. Both of these phenotypes indicate that initiation is either inefficient or diminished by the E to G mutation. This step requires the reorganization of actin into parallel filaments that continue to grow through the action of actin polymerases such as VASP or formins. A reduction in motor recruitment to the cortex via actin binding (as is the case for DdMyo7) or a decreased ability to interact with F-actin, or both, would result in decreased assembly of filopodia initiation foci that, in turn, would result in fewer filopodia.

Filopodia extension is driven by the activity of actin polymerases, such as VASP or formins, at the tip. Recent work has suggested that tip-localized Myo10 also has a role in elongation by reducing membrane tension to facilitate the addition of G-actin monomers (17, 18). Delivery of additional Myo10 motors by intrafilopodial motility (30) may ensure maintenance of sufficient amounts of motor at the tip to contribute to ongoing elongation. If so, then one would predict that mutation of the F-actin binding interface of Myo10 would have a significant impact on the length of Myo10-generated filopodia. The impact of the E to G mutation on filopodial length depends on the myosin and, for Myo10, on the cell type. In the case of DdMyo7-E456G, filopodia are only 18% shorter, indicating that this filopodial myosin does not have a major role in elongation. In contrast, the impact of the Myo10-E508G mutant on filopodia length varies, from significant reduction to actually increasing elongation (Table 2, Supp Table 1). Myo10-E508G resulted in a slight lengthening of filopodia in COS-7 cells (12%, Table 2), while it decreased the length of filopodia in U2-OS and HeLa cells either modestly or significantly, ∼ by 20% and 50%, respectively (Supp Table 1). The basis for the inconsistent effect of the *jd* mutation of mammalian cells filopodial length is unclear but it could be attributed to significant differences in the average length of filopodia that each cell line makes. The COS-7 cells extend the shortest filopodia (2.04 ±0.02 µm), U2-OS filopodia are somewhat longer (2.79 ± 0.03 µm) and HeLa cell filopodia are significantly longer (4.71± 0.04 µm) (Table 2, Supp Table 1). Notably, the HeLa cell filopodia length is impacted the most by Myo10-508G mutation, possibly due to a combination of much slower intrafilopodial transport down the long filopodia (Fig 5, Table 3) and compromised force generation at the tip due to reduced actin binding. It is also possible that the longer filopodia experience greater membrane tension at the tip. If there are fewer, less functional motors localized there, then there would be less force generated than needed to reduce this tension (Table 2, Supp Table 1). Alternatively, fewer Myo10 motors at the tip that are not fully functional could impact the extension cycle of filopodia that depends on substrate adhesion at the tip (28, 31). This could be reduced if levels of activated β-integrin at the tip were diminished (32) and may result in destabilization of the filopodium, impairing elongation. The inconsistent impact of the Myo10 *jd* mutant on filopodia length contrasts with the consistent reduction of filopodia number observed in all cell types (Table 2, Supp Table 1). These results show that the major function of Myo10 is to organize filopodial initiation sites essential for the formation of a filopodium and suggests that it makes less significant contributions to their elongation.

### A conservative substitution in the HTH impacts myosin function

Alignment of a wide range of myosin motor domain sequences showed that the HTH *jd* residue is predominantly an E but it is a D in a small number of myosins (Fig 1C). While the overall charge is conserved, the two amino acids differ by a single carbon, thus the length of the two residues is slightly different. Given their close similarity, it was initially considered that substitution of the E with a D would not have a major impact on the function of the filopodial myosins. However, introducing an D into the HTH *jd* position notably decreased the ability of DdMyo7 and Myo10 to generate filopodia. In the case of the DdMyo7-E456D, a smaller fraction of cells make filopodia compared to the WT (44 % versus 77%), with cells making 50% fewer filopodia/cell that were slightly shorter (Figs 2B, 3A, Table 1). Tip intensity was also significantly lower (Fig 3B, Table 1). Similarly, the in vivo activity of the Myo10-E508D mutant was also impaired by the conservative substitution, but this depended on the cell line. Filopodia production in COS-7 cells was reduced by 40% (Fig 4B,Table 2) but little or no impact on filopodia number was seen in either U2-OS or HeLa cells (Supp Fig 1A, C, Supp Table 1). In spite of slower intrafilopodial speed (Fig 6, Table 3), Myo10-E508D has a modest effect on tip intensity but no effect on the length of filopodia in U2-OS and HeLa cells. The mutation actually promoted a small increase the length of COS-7 filopodia (Fig 4B, Table 2, Supp Fig 1B, D, Supp Table 1). A major difference between the mammalian cell lines used in this study is that COS-7 cells make few filopodia while both U2-OS and HeLa cells make abundant filopodia. The basis for this difference is not known, but variations in the amounts and/or activity of filopodial actin polymerases and cross-linkers could compensate for the altered Myo10 activity. Overall, the results are consistent with the DdMyo7-E456D and Myo10-E508D mutations affecting their initiation activity. These finding establish that the even a modest change in the HTH region of the myosin surface that interacts with F-actin disrupts filopodial myosin function. The results also highlights how a seemingly minor change in the critical actin binding HTH region of the myosin motor can notably alter motor functionality in vivo.

### Conservation of filopodial myosin activity

DdMyo7 and Myo10 are evolutionarily distant filopodia myosins that are functional homologues that have similar roles. They are both tightly localized at the filopodia tip and are essential for filopodia formation (4, 5, 33). While they are both MF family members, their motor properties are different, the amoeboid motor is slow (∼ 40 nm/sec) while the mammalian one is significantly faster (∼ 600 nm/sec) (12, 13). The differences in motor activity would suggest that they play distinct roles in filopodia. Interestingly, introduction of the ‘*jd*’ mutation into the HTH region of these myosins results in quite similar phenotypes. The shared reduction of filopodia number generated when the jd mutation is introduced into either DdMyo7 or Myo10 implicates both myosins as major contributors to process of initiation. These myosins are likely to play a role in the initial alignment of parallel actin filaments and in maintaining a link between them as during assembly of the the nascent filopodial actin bundle. While loss of either DdMyo7 or Myo10 has some effects on filopodia elongation, these myosins do not appear to have fundamentally important roles in this process in all cell types indicating that their roles can vary in this respect. In spite of significant differences in motor activity and how they are recruited to the cortex, the shared requirement for DdMyo7 and Myo10 in filopodia initiation reveals strong evolutionary conservation of filopodial myosin function.

## Materials and Methods

### Protein alignments

Select motor domain sequences from representative myosin family members across phylogeny were downloaded from Cymobase (cymobase.org) (34). Note that recently identified classes have fewer representative sequences and are under-represented in the alignment. The full motor domains sequences were aligned using the Clustal Omega server hosted by the European Bioinformatics Institute (https://www.ebi.ac.uk/jdispatcher/msa/clustalo). The alignment file was opened in Jalview (https://www.jalview.org) (35), edited to only include the HTH region and the consensus visualized using WEBLOGO (https://weblogo.berkeley.edu) (36).

### Cell lines, cell maintenance, and transformations

*Dictyostelium* wild-type control (G1-21) or *myo7* null Ax3 cells (HTD17-1) (5) were cultured in HL5 media (Formedium) supplemented with 60 U/mL penicillin G and 60 ug/mL streptomycin sulfate (Sigma-Aldrich). Transgenic lines were generated by electroporation (27, 37) and selected with 10-20 µg/ml G418 (Fisher Scientific) or 50 ug/ml Hygromycin (Gold Biotechnology) and screened for expression by microscopy and western blotting.

Expression of fusion proteins was verified by western blotting using anti-Myo7 (UMN87) with anti-MyoB used as a loading control (38). Bands were detected using fluorescent secondary antibodies (LiCor) imaged with the LiCor Odyssey.

COS-7 (American Type Culture Collection, CRL-1651), U2-OS (American Type Culture Collection, HTB-96), and HeLa (American Type Culture Collection, CCL-2) cells were maintained at 37°C with 5% CO2 in either Dulbecco’s modified Eagle’s medium (COS-7 and HeLa; Thermo Fisher Scientific) or McCoy’s 5A modified medium (U2-OS; Thermo Fisher Scientific). Both media types were supplemented with 10% fetal bovine serum (FBS; Thermo Fisher Scientific) and 1% penicillin streptomycin (Gibco). For transfections, 0.25-1 µg plasmid DNA was combined with lipofectamine 2000 (Thermo Fisher Scientific) in Opti-MEM I reduced serum media (Thermo Fisher Scientific) and allowed to incubate for 5-10 minutes before adding the mixture dropwise to cells at 60-80% confluency plated in a 12-well plate in antibiotic-free media. For confocal microscopy experiments, cells were allowed to incubate in DNA for 24 hours before trypsinizing and replating on fibronectin for imaging. For TIRF microscopy experiments, cells were transfected and replated within 6-8 hours to ensure the low levels of expression necessary for single-molecule imaging.

Western blotting was used to verify expression of fusion proteins. eGFP Myo10 was detected with anti-human Myo10 (Santa Cruz Biotechnology sc-166720 1:500) with anti-human vinculin (Proteintech 66305-1-Ig, 1:10,000) or anti-beta tubulin (Proteintech 66240-1-Ig, 1:20,000) used as a loading control. Bands were detected using fluorescent secondary antibodies (LiCor) imaged with the LiCor Odyssey.

### Generation of expression plasmids

A combination of PCR- and restriction enzyme-based cloning was used to create Ddisc wild type and mutant mNeon-DdMyo7 expression plasmids in an mNeon-pDM304 expression plasmid (27, 39). Unless noted, cloning enzymes were obtained from New England Biolabs. A motor domain subclone (pDTi324) was generated by PCR cloning of the region coding amino acids 1 - mutagenized using Q5 mutagenesis (New England Biolabs) to introduce either the E456G or E456D codons and then restriction enzyme cloning was used to exchange the 5’ region of the gene carrying the E456G or E456D motor mutations into the wild type pDTi516 mNeon-DdMyo7 expression plasmid (27), creating pDTi531 and pDTi532 respectively.

The mammalian eGFP-Myo10 expression plasmid was obtained from Addgene (#47609; a gift from Emanuel Strehler, (40)) and mutations introduced by mutagenic PCR (CloneAmp, Takara) followed by KLD mix ligation (New England Biolabs). The sequences of each PCR-generated clone were confirmed by Sanger sequencing (GeneWiz/Azenta) or whole plasmid sequencing (Plasmidsaurus).

### Microscopy and imaging experiments

Microscopy of living Ddisc cells was carried out as previously described (7). Briefly, the cells were adhered to glass bottom imaging dishes (CellVis, D35-10-1.5-N) and starved for 1-2 hours in nutrient free buffer (SB, 16.8 mM phosphate, pH 6.4) at 19-21°C. Cells were plated at a density of 5 x 10^5^ per ml were imaged on a Nikon Ti2-E microscope equipped with a Crest Optics X-Light V3 spinning disk and captured with a Hamamatsu ORCA-Fusion BT CMOS camera, 200 µm stagetop Piezo Z for High-Speed Z-stack Acquisition, and an OKO stagetop incubator. Samples were illuminated with Lumencor CELESTA light engine 800 mW lasers (488 nm) with GFP or DS red filters (High Signal to Noise BL Series, Nikon) for 488-nm excitation (final 0.1099 µm pixel size). Imaging was performed using a 60 x Plan Apo oil-immersion objective (NA 1.4), 4-6 Z sections of 0.3 µm were taken with 50-100 ms exposure with 30-60% laser power over 10-20 s. Ten fields of view were collected from each imaging dish, with 2-20 cells per field of view. All data sets represent cells from at least three independent experiments and two independently transformed cell lines.

Cells were trypsinized the day following transfection and replated on 24-well glass bottom plates with #1.5 coverslips (CellVis) coated in 5µg/mL fibronectin (Sigma Aldrich) for confocal imaging. Cells were plated at a low density to prevent filopodia or cell edge overlap and were allowed to adhere for at least two hours before replacing media with Fluorobrite DMEM (Thermo Fisher Scientific) supplemented with 10% FBS. The same microscope, spinning disk, camera, objective, lasers, and filters as described for Ddisc imaging were used for mammalian cell imaging. Five z-stacks of 0.3 µm each were taken with a 1s exposure at 60% laser power. Imaging was performed at 37°C maintained by an OKO stagetop incubator.

### Image and data analysis

Filopodia analysis and cortex to cell ratio analysis was done on maximum intensity projections using a custom automated deep learning platform ‘filoTips’ (26). Cells not expressing transgenic proteins were excluded from the analysis. Values from filoTips were analyzed and graphed using GraphPad Prism v. 10. Data points deemed definite outliers (0.1%) by Rout method were excluded, then one-way ANOVA analysis with post hoc Dunnett’s multiple comparison between wild-type control and each mutant was used to compare groups. Kymographs were generated by manually tracing filopodia lengths and using the reslice command in ImageJ (41). Velocity measurements were obtained by tracing Myo10 tracks on kymographs and calculating the velocity via the angle of the line then plotted in R and fit to a Gaussian curve. Statistical tests were calculated and experimental means with error bars are SEM (standard error of the mean) are shown on graphs to demonstrate experimental variability. Significant differences were compared to the control or between mutants expressed in the same genetic background, unless noted.

## ACKNOWLEDGEMENTS

The project was supported by a grant from the National Institute of General Medical Sciences (R01GM122917 to MAT) and the Agence Nationale de la Recherche (ANR-19-CE11-0015-02, ANR-24-CE13-1876-03 to AH). The NikonTi2-E and Crest V3 spinning disc confocal and Licor Imager were purchased with Equipment Supplement awards (3R01GM122917-02S1, 3R01GM122917-03S2 and 3R01GM122917-04S1, respectively). The OKO Stagetop Incubator was purchased with funds from a Grant-In-Aid of Research, Artistry and Scholarship (#551650) from the University of Minnesota and matched funding from the Department of Genetics, Cell Biology and Development. Part of the work reported here was supported by the resources and staff at the University of Minnesota University Imaging Centers (UIC, RRID SCR_020997).

## SUPPLEMENTARY FIGURES AND TABLE

**Supplemental Figure 1.**
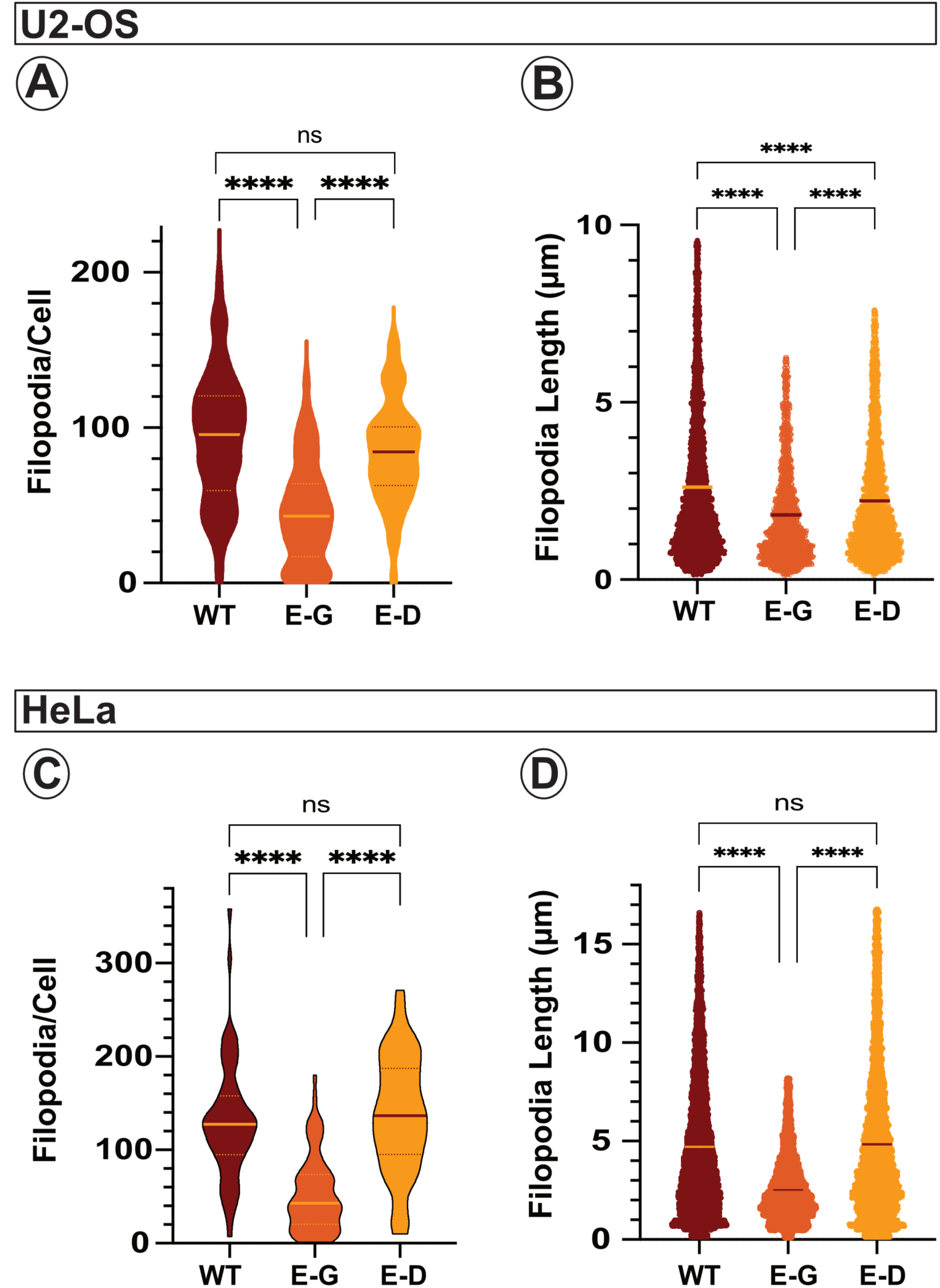
The Myo10 ‘*jd*’ mutation reduces filopodia formation in both U2-OS and HeLa cells. Quantitative analysis of filopodia formation in mammalian cell lines expressing either the wild type eGFP-Myo10 (WT), E508G (E-G) or E508D (E-D) using filoTips. Top - U2-OS cells, **A.** Filopodia per cell. **B.** Filopodia length. Bottom - HeLa cells. **C.** Filopodia per cell. **D.** Filopodia length. **A, C** The mean is indicated with a thick bar on the plot for each line and thinner lines indicate the quartiles. **B, D** The mean is indicated with a thick bar on the plot for each line. Shown are the results from two independent cell lines for each myosin, ** p < 0.002, **** p < 0.0001 ns = not significant

**Supplemental Figure 2.**
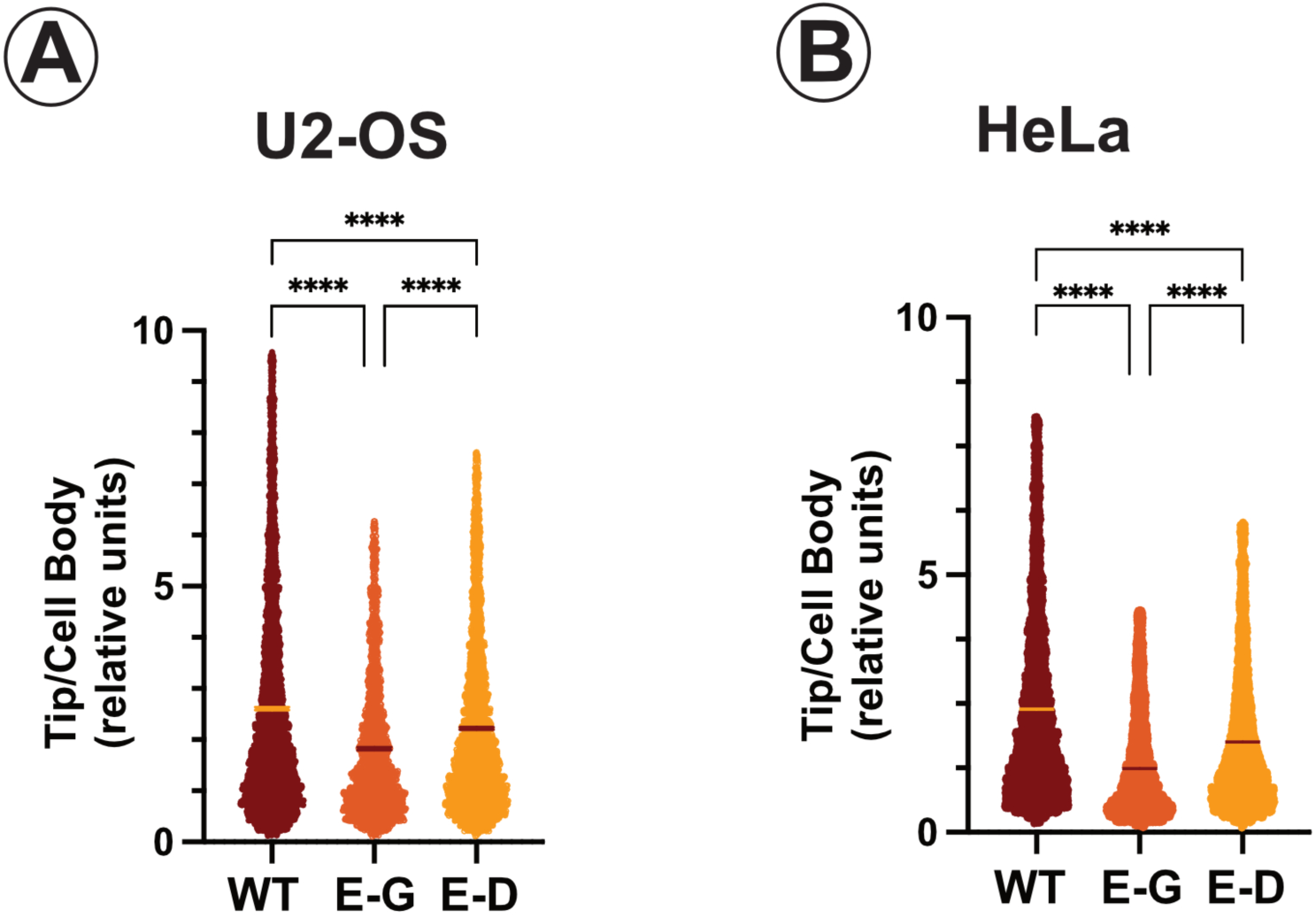
Varied reduction of filopodia tip intensity in the Myo10 ‘*jd*’ mutation in U2-OS and HeLa cells. Quantitative analysis of the tip intensity of mammalian cells expressing either the wild type eGFP-Myo10 (WT), E508G (E-G) or E508D (E-D) using filoTips. **A.** U2-OS. **B.** HeLa. Shown are the results from the analysis of two independent cell lines for each myosin, the mean is indicated with a thick bar on the plot for each line. **** p < 0.0001

**Supplementary Table 1.**
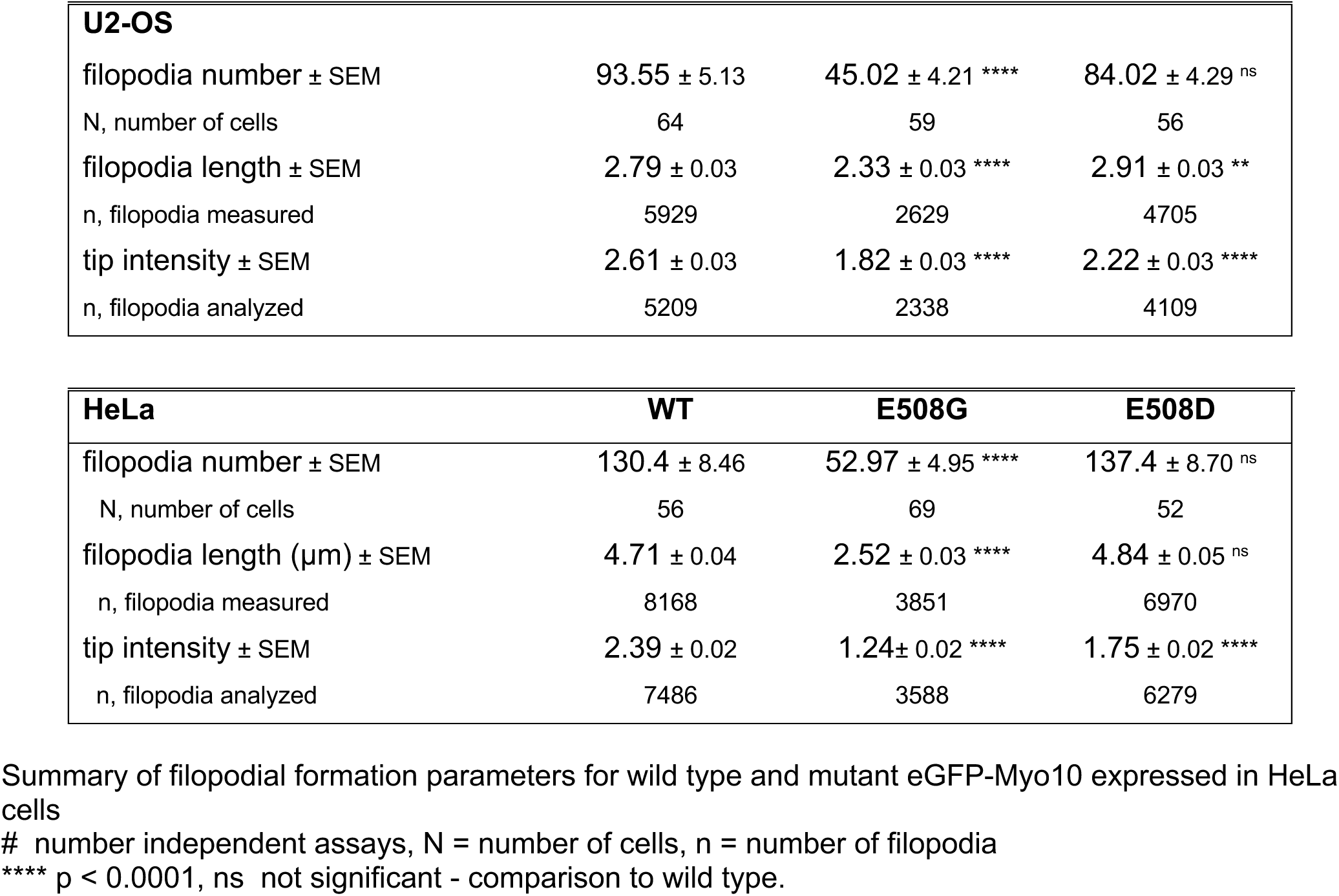
Impact of HTH mutants on Myo10 filopodial function in U2-OS and HeLa Cells.

